# Rac2 Hyperactivity Drives Neutrophil Degranulation, Myeloperoxidase Deficiency, and Lymphopenia

**DOI:** 10.64898/2026.05.12.723629

**Authors:** H. Hanson, M. Rodriguez, E. Kugelmann, M. Malafei, M. Boe, D. J. Montell

## Abstract

Patients with a dominant mutation in the Rho GTPase RAC2, RAC2^E62K^, which hyperactivates the protein, suffer from a combined immunodeficiency characterized by recurrent bacterial and fungal infections and severe T cell lymphopenia. Patient neutrophils have elevated F-actin and superoxide production yet fail to control growth of *S. aureus*, and the mechanism underlying this killing defect is unknown. Here we report that hyperactive Rac2 primes neutrophils for primary granule degranulation, potentially depleting myeloperoxidase (MPO) needed for intraphagosomal microbial killing. Using a Rac2^+/E62K^ mouse model, we show that mature bone marrow neutrophils have decreased side scatter, elevated surface CD63, and reduced intracellular MPO. Interestingly, bone marrow architecture and neutrophil development in the mice are normal. Rac2^+/E62K^ neutrophils are hyperactivated, with increased CD11b expression, cell spreading, and bioparticle phagocytosis. In the spleen, Rac2^+/E62K^ mice display extramedullary granulopoiesis and an accumulation of degranulating neutrophils. Splenic T cells, but not B cells, show elevated surface phosphatidylserine, an “eat me” signal that sensitizes them to phagocytic clearance and provides a candidate mechanism for the selective T cell lymphopenia. Together these findings suggest that hyperactive Rac2 compromises antimicrobial neutrophil function and drives selective T cell clearance in the spleen.

## Introduction

Patients with a dominant hyperactivating mutation in the Rho GTPase RAC2, RAC2^+/E62K^, present with neutrophil dysfunction and profound lymphopenia (Hsu et al., 2019; Smits et al., 2020). Patients are susceptible to S. *aureus*, *Aspergillus*, and recurrent sinopulmonary bacterial infections, implicating a defect in neutrophil oxidative killing, despite having normal numbers of circulating neutrophils that produce (Hampton et al., 1996; Aratani et al., 2002; Smits et al., 2020). RAC2^+/E62K^ neutrophils produce excessive superoxide and have increased F-actin, macropinocytosis, and impaired chemotaxis, all of which are consistent with known functions of Rac (Hsu et al., 2019; Smits et al., 2020). Smits et al. also reported that neutrophils cannot stop S. *aureus* growth, but the mechanism behind this defect was not identified and is surprising given the elevated superoxide production.

RAC2 expression is restricted to hematopoietic cells and drives cytoskeletal remodeling through actin polymerization (Burridge and Wennerberg, 2004). The E62K mutation enhances RAC2 activity by altering interactions with regulators such as GTPase activating proteins, thereby increasing the time spent in a GTP bound state (Hsu et al., 2019; Arrington et al., 2020). RAC2 is essential for primary antimicrobial effector functions in neutrophils including phagocytosis and the release of extracellular traps (NETosis) (Torfs et al., 2026; Lim et al., 2011). It is also crucial for primary (azurophilic) degranulation (Abdel-Latif et al., 2004). RAC2-driven actin polymerization brings granules to the cortex, while local F-actin disassembly at the membrane is required for granule fusion and content release (Jog et al., 2007; Mitchell et al., 2008). Primary neutrophil degranulation is a tightly regulated process in which granules fuse with the membrane and release antimicrobial agents extracellularly (Lacy, 2006). Granules are loaded with cytotoxic agents, digestive, and antimicrobial enzymes such as elastase and myeloperoxidase (MPO) (Othman et al., 2022). MPO reacts with hydrogen peroxide and chloride to generate hypochlorous acid (HOCl), a potent oxidant required for killing phagocytosed pathogens (Klebanoff 1968, Klebanoff 2005).

Granules are synthesized early in development, at the promyelocyte stage, but MPO mRNA levels decline and are absent in fully mature neutrophils, so they are a non-renewable resource (Van der Veen, 2009). We hypothesized that hyperactive RAC2^+/E62K^ might lead to premature azurophilic degranulation, depleting intracellular myeloperoxidase stores and other antimicrobial agents in primary granules necessary for microbial killing.

RAC2^+/E62K^ patients also suffer from severe T cell lymphopenia. Rac2 is expressed and important for mediating cytoskeletal changes during T cell development and activation (Dumont et al., 2009; Yu et al., 2001). However, bone marrow and thymic T cell development in a murine Rac2^+/E62K^ model is normal, leaving the peripheral T lymphopenia unexplained (Hsu et al., 2019; Smits et al., 2020; Mishra et al., 2023). Mishra et al. showed that Rac2^+/E62K^ bone marrow derived macrophages are hyperphagocytic and that Rac2^+/E62K^ T cells are sensitized to engulfment compared to wild type T cells. However, the mechanisms causing T cell sensitivity to engulfment remain unknown.

The Rac2 protein differs by only two amino acids in mice and humans (Didsbury et al., 1989). A Rac2^+/E62K^ heterozygous mouse model recapitulates human patient phenotypes including increased neutrophil F-actin, enhanced superoxide production, and T cell lymphopenia (Hsu et al., 2019). Here, we used the mouse model to investigate neutrophil development and function to identify additional mechanisms that might contribute to the combined immunodeficiency. We show that the E62K mutation primes neutrophils for degranulation. Neutrophils show hyperactivated phenotypes including increased cell spreading, CD11b expression, and enhanced phagocytosis of bioparticles. This priming towards degranulation and subsequent MPO defect may also explain the discrepancy between increased internalization but decreased killing power of live S. *aureus* previously published. Rac2^+/E62K^ mice have severe splenic extramedullary granulopoiesis and an accumulation of degranulating neutrophils. Splenic T cells have significantly elevated PS exposure consistent with enhanced phagocytic clearance, providing a mechanism for T cell lymphopenia. Together these results suggest mechanistic explanations that likely contribute to RAC2^+/E62K^ patients’ increased susceptibility to bacterial infections and T cell lymphopenia.

## Results

### Rac2^+/E62K^ neutrophils develop normally but display markers of primary granule degranulation

RAC2^+/E62K^ patients present with neutrophil dysfunction despite relatively normal circulating neutrophil counts, raising the question of whether the mutation impairs neutrophil function, development, or both (Smits et al. 2020). RAC2 expression is moderate in hematopoietic stems cells, decreases during the myelocyte stage, and reaches maximum expression in mature neutrophils (CZ CELLxGENE Discover). To investigate, we examined bone marrow neutrophils, isolated from the Rac2^+/E62K^ mice, across a panel of cytoskeletal, functional, and developmental readouts. We first measured side scatter across mouse Rac2^+/E62K^ myeloid populations by flow cytometry. Mature (LY6G-hi/ CD11b-hi/CXCR2-hi) Rac2^+/E62K^ neutrophils in the bone marrow have significantly decreased side scatter (Fig. 1A). However, the side scatter of immature neutrophils and preNeu populations do not significantly differ, demonstrating that the phenotype is specific to cells later in development, when Rac2 expression is highest. This result suggests that mature Rac2^+/E62K^ neutrophils have reduced internal complexity or granularity.

**Figure 1.**
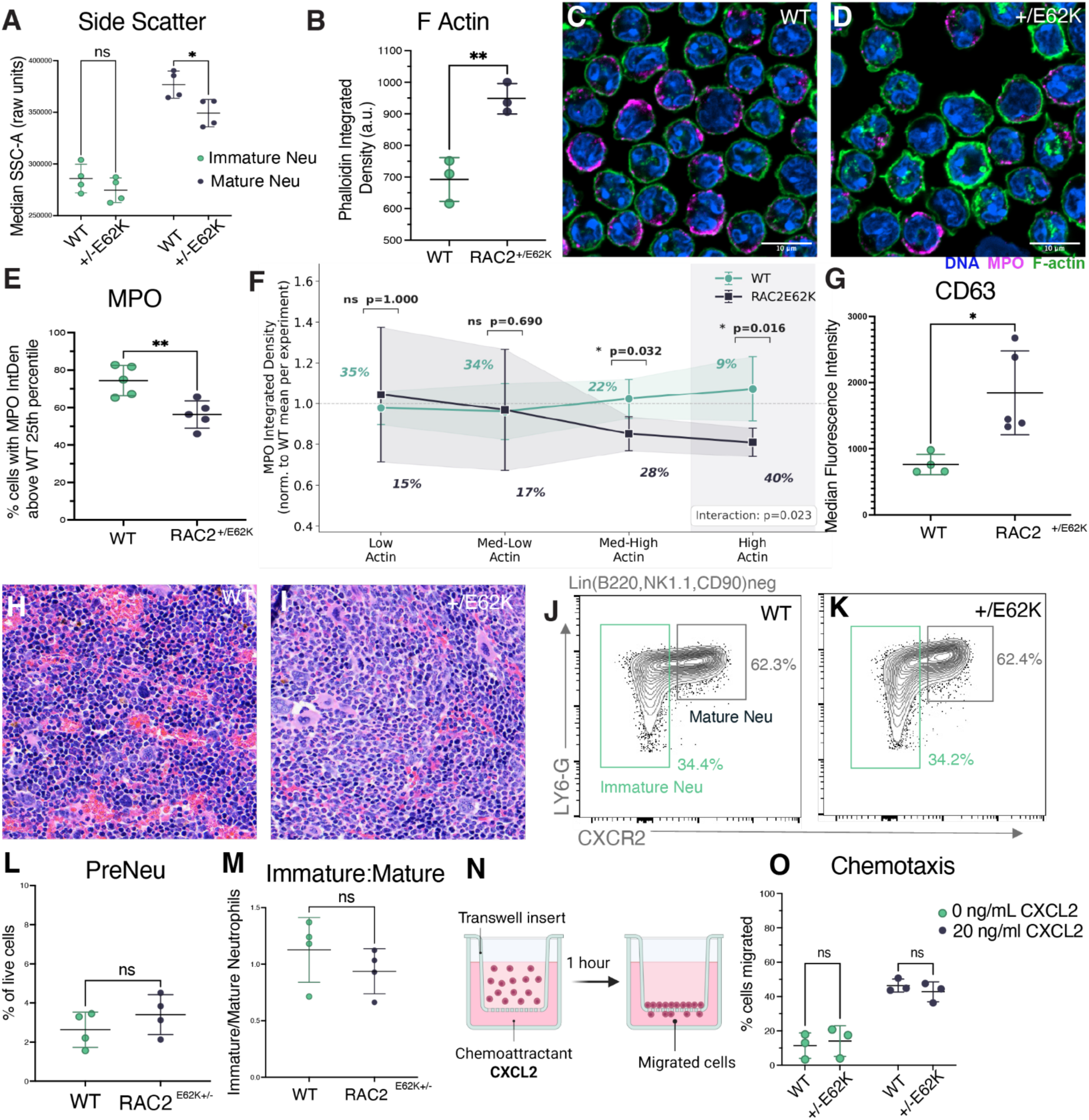
Rac2^+/E62K^ bone marrow neutrophils show degranulation and myeloperoxidase deficiency. (A) Median side scatter area (SSC-A) of immature and mature bone marrow neutrophil populations measured by flow cytometry. n=4 mice per genotype. Two-way ANOVA with Tukey’s post-hoc test (factors: genotype, maturation state). (B) F-actin content of bone marrow neutrophils measured as average phalloidin integrated density per cell from confocal images. n=3 mice per genotype. Unpaired t-test. (C–D) Representative confocal images of bone marrow neutrophils from WT (C) and Rac2^+/E62K^ (D) mice stained with phalloidin-488 (F-actin, green), anti-MPO antibody (magenta), and Hoechst (nuclei, blue). Scale bar = 10 µm. (E) Percentage of bone marrow neutrophils with MPO integrated density above the WT 25th percentile threshold, calculated within each experiment to control for batch variability. n=5 mice per genotype. Unpaired t-test. (F) MPO integrated density normalized to the WT mean per experiment, plotted across F-actin quartile bins defined on the pooled normalized F-actin distribution. Percentages indicate the fraction of each genotype’s cells within each bin. n=5 mice per genotype. Per-bin comparisons by Mann-Whitney U test on per-mouse bin means; interaction p-value from OLS regression with genotype, F-actin bin, and their interaction as predictors. (G) Median fluorescence intensity (MFI) of CD63 expression on mature bone marrow neutrophils by flow cytometry. n=5 mice per genotype. Unpaired t-test. (H–I) Hematoxylin and eosin staining of femurs from WT (H) and Rac2^+/E62K^ (I) mice at 40× magnification. (J–K) Representative flow cytometry contour plots of Lin⁻ (B220/NK1.1/CD90.2-negative) bone marrow cells from WT (J) and Rac2^+/E62K^ (K) mice showing LY6G versus CXCR2 expression to distinguish immature (teal gate) from mature (grey gate) neutrophils. Numbers indicate percentage of parent gate. (L) PreNeu population as a percentage of live bone marrow cells by flow cytometry. n=4 mice per genotype. Unpaired t-test. (M) Ratio of immature to mature bone marrow neutrophils as defined in (J–K). n=4 mice per genotype. Unpaired t-test. (N) Schematic of the transwell migration assay. (O) Quantification of bone marrow neutrophil chemotaxis toward 0 or 20 ng/mL CXCL2 over 1 hour. n=3 mice per genotype. Two-way ANOVA with Tukey’s post-hoc test (factors: genotype, CXCL2 concentration). Each dot represents one mouse, except panel I which shows the average of 5 mice. Imaging panels (B, E, F): cells gated to 40–120 µm² area. Data are mean ± SD; *p<0.05, **p<0.01, ***p<0.001, ****p<0.0001; ns = not significant

Published characterization of Rac2^+/E62K^ patient neutrophils described elevated F-actin (Hsu et al. 2019). To determine if this was recapitulated in mouse bone marrow neutrophils, we compared F-actin levels using confocal microscopy and phalloidin staining. We found that F-actin levels are enhanced in Rac2^+/E62K^ neutrophils prior to egress from the bone marrow (Fig. 1B).

To further investigate granularity in bone marrow neutrophils, we measured total MPO, an enzyme found in primary granules (Fig. 1C, 1D). Amongst wild type neutrophils, MPO levels vary (Fig. 1E), so we binned cells into quartiles. A significantly lower percentage of Rac2^+/E62K^ neutrophils have myeloperoxidase levels above the 25^th^ percentile (p=0.005, unpaired t-test, n=5 mice per genotype). A subset of Rac2^+/E62K^ neutrophils retained MPO levels comparable to WT controls, arguing against a protein synthesis defect. Instead, MPO depletion appeared to scale with phalloidin intensity, suggesting that cells might begin with equivalent MPO stores and then, as Rac expression and activity increase and F-actin accumulates, MPO stores might be lost.

To quantify the relationship between F-Actin and MPO, we established low, medium-low, medium-high, and high actin quartiles based on phalloidin intensity. We found that at low phalloidin intensities, the MPO levels did not differ between WT and Rac2^+/E62K^ (Fig. 1F). However, at higher F-actin levels, there was a significant decrease in Rac2^+/E62K^ MPO (med-high p= 0.032, high p=0.016). Rac2^+/E62K^ neutrophils were depleted of MPO in proportion to their F-actin intensity, suggesting that elevated Rac2 activity might sensitize cells to granule release. The relationship between F-actin level and MPO content differed between genotypes. We conclude that Rac2^+/E62K^ neutrophils show progressive MPO depletion as F-actin increases, which could be due to altered development or degranulation.

To distinguish between the two possible mechanisms of MPO loss, we examined cell surface CD63, a marker of degranulation. CD63 is localized to the primary granule membrane and appears extracellularly only after granule fusion with the plasma membrane during degranulation (Niessen and Verhoeven, 1992; Källquist et al., 2008). To test whether decreased MPO reflects release of degranulation, we measured extracellular CD63 expression by flow cytometry (Fig. 1G and Fig. S1L). Rac2^+/E62K^ neutrophils show significantly increased CD63, suggesting that the Rac2^+/E62K^ mutation may prime neutrophils for degranulation.

To investigate the possibility that decreased MPO content reflects altered neutrophil development, we compared the bone marrow from WT and Rac2^+/E62K^ mice. Hematoxylin and eosin staining of WT and Rac2^+/E62K^ femurs revealed no gross difference in architecture, with comparable organization and cellularization at low and higher magnification (Fig. 1H and I and Fig. S1A and B,). To assess neutrophil development quantitatively, we performed flow cytometry immunophenotyping using LY6G and CXCR2 to differentiate immature and mature neutrophil populations (Fig. 1J and K and Fig. S1C-J). The percentage of PreNeu or immature neutrophils as a total of live cells did not differ significantly (Fig. 1L). The ratio of immature to mature neutrophils also did not differ (Fig. 1M). Similarly, Smits et al. 2020 found no bone marrow abnormalities in a post mortem evaluation of a human RAC2^+/E62K^ patient.

Hsu et al. described impaired fMLF-directed chemotaxis of circulating neutrophils from RAC2^+/E62K^ patients. We wondered if a neutrophil chemotaxis defect might impair egress from the bone marrow. CXCL2, which is a major chemoattractant involved in bone marrow egress (Eash et al., 2010), so we performed a transwell migration assay to measure neutrophil chemotaxis to CXCL2 (Fig. 1F). We observed no significant difference in chemotaxis between WT and Rac2^+/E62K^ neutrophils either in the absence or presence of CXCL2 (Fig. 1G). This suggests that developing neutrophils should be able to exit the bone marrow normally. Together, these data indicate that Rac2^+/E62K^ bone marrow neutrophil development proceeds normally, demonstrating that the functional phenotypes described below reflect the effect of increased Rac activity rather than indirect effects on the maturation status of individual cells.

### Rac2^+/E62K^ neutrophils are hyperactivated but have impaired microbicidal activity

To determine whether mutant neutrophils are functionally hyperactivated, we measured cell spreading on fibronectin with and without GM-CSF stimulation. Without stimulation, Rac2^+/E62K^ neutrophils exhibited enhanced spreading compared to wild type (Fig. 2A). Following GM-CSF stimulation, although the trend was the same, the differences did not reach statistical significance, suggesting that the mutation mimicked low-level stimulation, consistent with the priming model.

**Figure 2.**
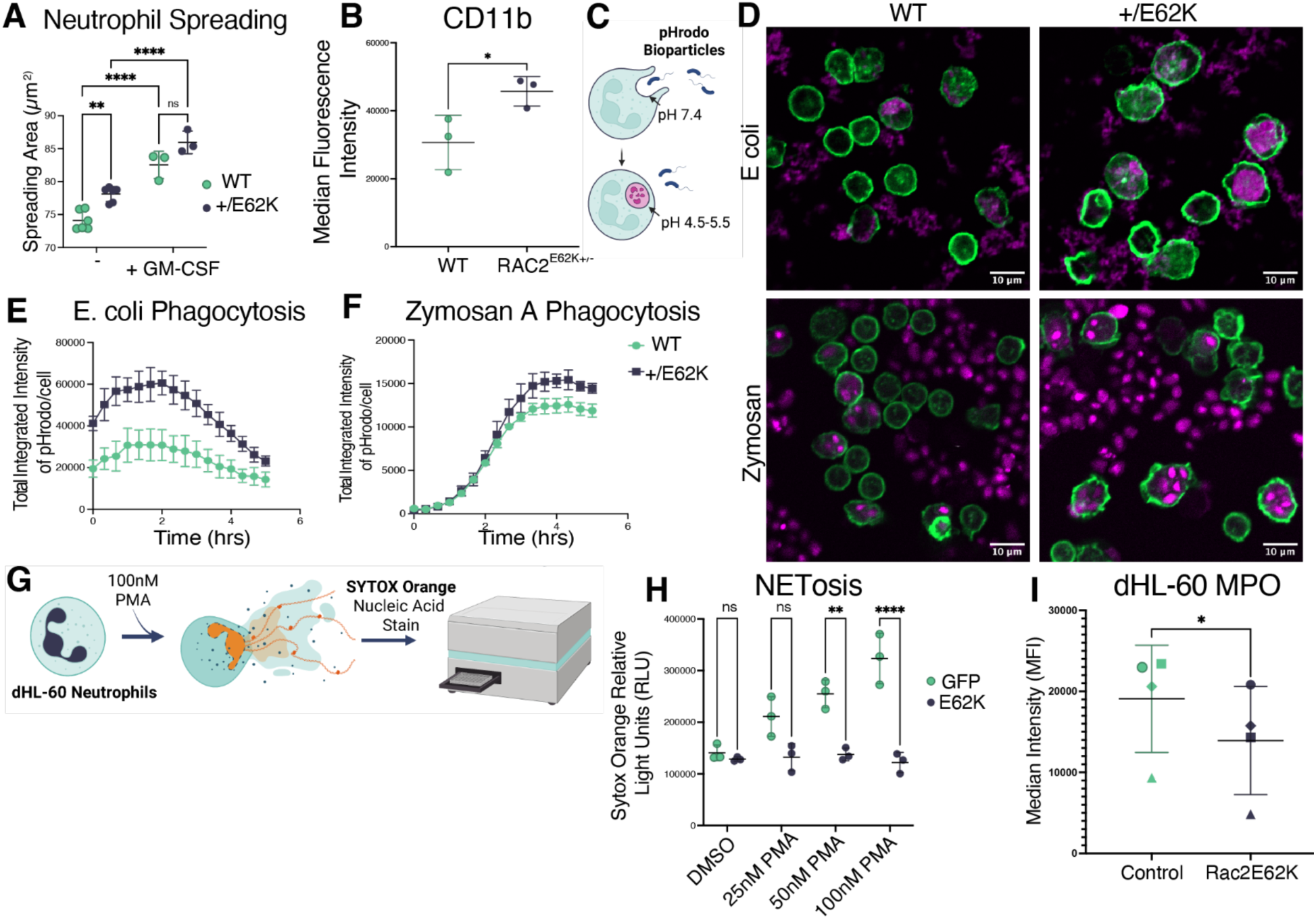
Rac2^+/E62K^ neutrophils are hyperactivated but may have impaired killing function. (A) Neutrophil spreading area on fibronectin-coated surfaces with and without GM-CSF stimulation after 2 hours. n=3 mice per genotype. Two-way ANOVA with Tukey’s post-hoc test (factors: genotype, GM-CSF stimulation). (B) Surface CD11b median fluorescence intensity (MFI) on bone marrow neutrophils by flow cytometry. n=3 mice per genotype. Unpaired t-test. (C) Schematic of the pHrodo bioparticle phagocytosis assay. (D) Representative confocal images of bone marrow neutrophils following coculture with pHrodo-labeled E. coli (upper) and Zymosan A (lower) bioparticles. F-actin in green, pHrodo signal in magenta. Scale bar = 10 µm. (E–F) Total pHrodo integrated intensity per cell over time for E. coli (E) and Zymosan A (F) bioparticles, imaged at 20-minute intervals for 5 hours. Lines represent mean per-cell pHrodo intensity per mouse. n=4 mice per genotype. Statistical comparisons performed on the area under the curve (AUC) per mouse by Mann-Whitney U test. (G) Schematic of the NETosis assay. (H) SYTOX Orange relative fluorescence units (RFU) quantifying NETosis in GFP control and RAC2^E62K^ dHL-60 cells at 0, 25, 50, and 100 nM PMA. n=3 biological replicates. Two-way ANOVA with Tukey’s post-hoc test (factors: genotype, PMA concentration). (I) Intracellular MPO MFI in GFP control and RAC2^E62K^ dHL-60 cells by flow cytometry. n=3 biological replicates. Unpaired t-test. Each dot represents one mouse (A, B, E, F) or one biological replicate (H, I). Data are mean ± SD; *p<0.05, **p<0.01, ***p<0.001, ****p<0.0001; ns = not significant.

CD11b (CR-3/Mac-1) is an integrin that drives cell adhesion and spreading (Hughes et al., 1992), expression of which is induced by GM-CSF (Van Pelt et al., 1996). Therefore, we asked whether differences in surface CD11b expression could explain the increased spreading of Rac2^+/E62K^ neutrophils. Indeed, hyperactive Rac increased CD11b expression in LY6G/CD11b+ neutrophils (Fig. 2B). Engagement of CD11b also activates signaling that drives cytoskeletal rearrangements and phagocytosis (Graham et al., 1989; Jongstra-Bilen et al., 2003). Therefore, we measured neutrophil phagocytosis of pHrodo E. *coli*, Zymosan A, and S. *aureus* bioparticles (Fig. 2C). Rac2^+/E62K^ neutrophils have enhanced bioparticle engulfment with intact acidification of internalized targets over time (Fig. 2D-F and Fig. S2D). For all three targets, the pHrodo intensity per cell is higher in the Rac2^+/E62K^ neutrophils. The phenotype is not restricted to a specific pathogen. This broad hyperphagocytic phenotype is important because while Smits et al. reported that patient neutrophils cannot stop the colonization of S. *aureus*, our results suggest that this may not be a phagocytosis defect, rather a killing defect.

NETosis is an MPO dependent response in which neutrophils decondense their nuclei and use it as an extracellular web to trap pathogens and stop the spread of infection (Torfs et al., 2026). Human, but not murine, neutrophils are dependent on MPO to undergo NETosis (Metzler et al., 2011). Therefore, we used the dHL-60 human neutrophil model to measure the effect of RAC2^E62K^ expression on NETosis. dHL-60 neutrophils expressing a doxycycline (Dox)-inducible RAC2^E62K^ were stimulated with 0, 25, 50, or 100nM of PMA, and NETs were measured by SYTOX orange fluorescence (Fig. 2G). At 5 hours, whereas wild type neutrophils exhibited increasing NETosis in response to increasing PMA concentration, RAC2^E62K^ neutrophils did not undergo significant NETosis at any concentration (Fig. 2H). We then asked if this NETosis defect is due to a degranulation driven MPO depletion. We measured MPO levels in dHL-60 neutrophils by flow cytometry and discovered that RAC2^E62K^ dHL-60 neutrophils have decreased MPO compared to the non-doxycycline-induced control (Fig. 2I). This result is consistent with our model that RAC2 expression drives degranulation. While decreased MPO may account for some of the NETosis defect, RAC2^+/E62K^ expressing cells are not entirely MPO deficient, making it unlikely that the MPO decrease fully explains the defect.

### Rac2^+/E62K^ Drives Splenic Extramedullary Granulopoiesis

Given the evidence for degranulation in bone marrow neutrophils, we next examined splenic neutrophil populations. The spleen acts as a reservoir for neutrophils for quick release into the blood stream (Jhunjhunwala et al., 2016). We performed H and E histology to understand how splenic compartments in the mice are impacted by Rac2^+/E62K^ neutrophilia (Hsu et al., 2019). In the white pulp, Rac2^+/E62K^ spleens have expanded germinal centers and expanded marginal zones that are multifocally coalescing (Fig. 3A). In the red pulp subcapsular regions, there are megakaryocytes, and increased numbers of myeloblast and band neutrophils, evidence of extramedullary hematopoiesis (Fig. 3B). Spleens from Rac2^+/E62K^ mice weighed significantly more than WT spleens (Fig. 3C).

**Figure 3.**
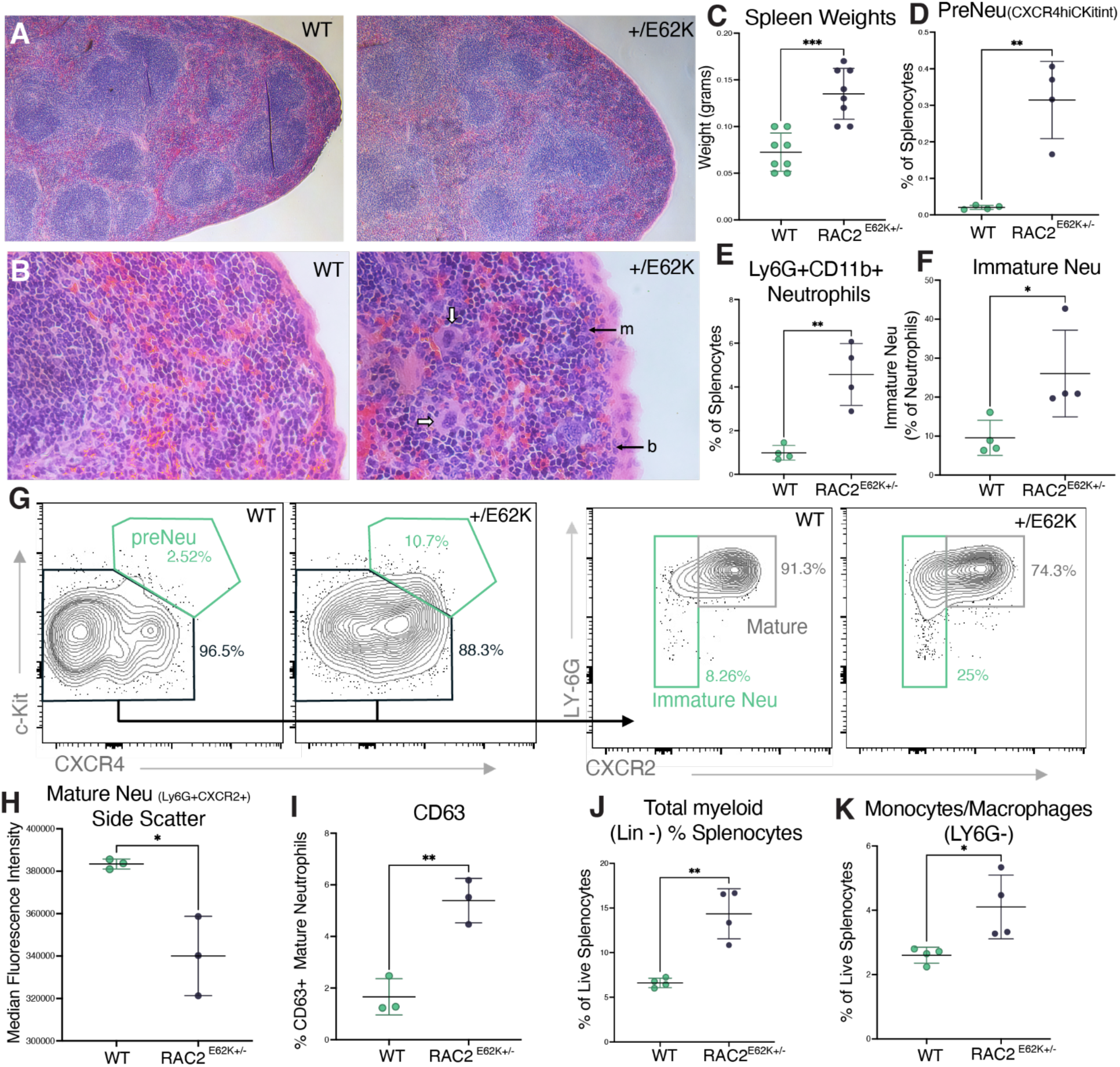
Rac2^+/E62K^ drives splenic extramedullary granulopoiesis in a mouse model. (A) Representative H&E staining of spleens from WT and Rac2^+/E62K^ mice at low magnification. (B) Higher-magnification H&E of splenic red pulp. Arrows indicate areas of extramedullary granulopoiesis in Rac2^+/E62K^ spleens. White arrows = megakaryocytes; m = myeloblast; b = band neutrophils. (C) Spleen weights. n=8 mice per genotype. Unpaired t-test. (D) PreNeu population (CXCR4-hi/c-Kit-int) as a percentage of total splenocytes by flow cytometry. n=4 mice per genotype. Unpaired t-test. (E) Ly6G⁺/CD11b⁺ mature neutrophils as a percentage of total splenocytes by flow cytometry. n=4 mice per genotype. Unpaired t-test. (F) Immature neutrophils as a percentage of total splenic neutrophils by flow cytometry. n=4 mice per genotype. Unpaired t-test. (G) Representative flow cytometry contour plots of Lin⁻ splenocytes. Left: c-Kit versus CXCR4 showing the preNeu population (teal gate, CXCR4-hi/c-Kit-int). Right: LY6G versus CXCR2 showing mature (grey gate, LY6G-hi/CXCR2-hi) and immature (teal gate, LY6G-lo/CXCR2-lo) neutrophils. Numbers indicate percentage of parent gate. (H) Median SSC-A of mature (Ly6G⁺/CXCR2⁺) splenic neutrophils by flow cytometry. n=4 mice per genotype. Unpaired t-test. (I) Percent CD63-positive mature splenic neutrophils by flow cytometry. n=4 mice per genotype. Unpaired t-test. (J) Total myeloid (Lin⁻) cells as a percentage of live splenocytes by flow cytometry. n=4 mice per genotype. Unpaired t-test. (K) Monocytes/macrophages (CD11b⁺/LY6G⁻) as a percentage of live splenocytes by flow cytometry. n=4 mice per genotype. Unpaired t-test. Each dot represents one mouse. Data are mean ± SD; *p<0.05, **p<0.01, ***p<0.001, ****p<0.0001; ns = not significant.

To confirm splenic extramedullary granulopoiesis, we used an immunophenotyping flow cytometry panel to evaluate the myeloid populations. We found a significant increase in the preNeu population in the spleen, which is normally restricted to the bone marrow (Fig. 3D). Overall, there was an increase in CD11b+ Ly6G+ neutrophil population in the Rac2^+/E62K^ spleens (Fig. 3E), and the immature neutrophil fraction (Fig. 3F). Figure 3G displays a representative gating strategy and population differences, suggesting extramedullary neutrophil development. Extramedullary granulopoiesis is consistent with compensation for defective neutrophil killing function and is broadly associated with an increase of immature neutrophils into circulation (Honda et al., 2016; Silvestre-Roig et al., 2026).

To investigate whether splenic neutrophil populations showed evidence of degranulation, we measured side scatter and CD63 expression. The side scatter of mature (Ly6G-hi/CD11b-hi/CXCR2-hi) Rac2^+/E62K^ neutrophils is significantly decreased compared to WT neutrophils (Fig. 3H), consistent with degranulation. The percentage of mature neutrophils positive for CD63 tripled in Rac2^+/E62K^ mice compared to wild type (Fig. 3I). Decreased side scatter and increased CD63 expression suggest that Rac2^+/E62K^ splenic neutrophils, like bone marrow neutrophils, are primed to degranulate.

We also measured the total myeloid and monocyte populations and found that there was an overall increase in the myeloid population in the Rac2^+/E62K^ spleen (Fig. 3J), consistent with published results from Hsu et al., as well as a significant increase in the monocyte/macrophage (CD11b+ LY6G-) population (Fig. 3K). The Rac2^+/E62K^ spleen is a myeloid rich environment populated with newly synthesized MPO rich precursor cells primed to degranulate as mature neutrophils.

### Rac2^+/E62K^ Splenic T cells Display Elevated PS Exposure

RAC2 is expressed and important for mediating cytoskeletal changes during T cell development and activation (Yu et al., 2001; Dumont et al., 2009). Given that hyperactive Rac activates neutrophils (this study) and macrophages (Mishra et al., 2023) we wondered if the lymphocytes show evidence of T cell hyperactivation. We measured F-actin by phalloidin staining, and used Incucyte imaging, and cell by cell analysis and found that Rac2^+/E62K^ T cells have increased F-actin compared to wild type (Fig. 4A-C). This result shows that the E62K mutation enhances F-actin content across multiple hematopoietic cell types.

**Figure 4.**
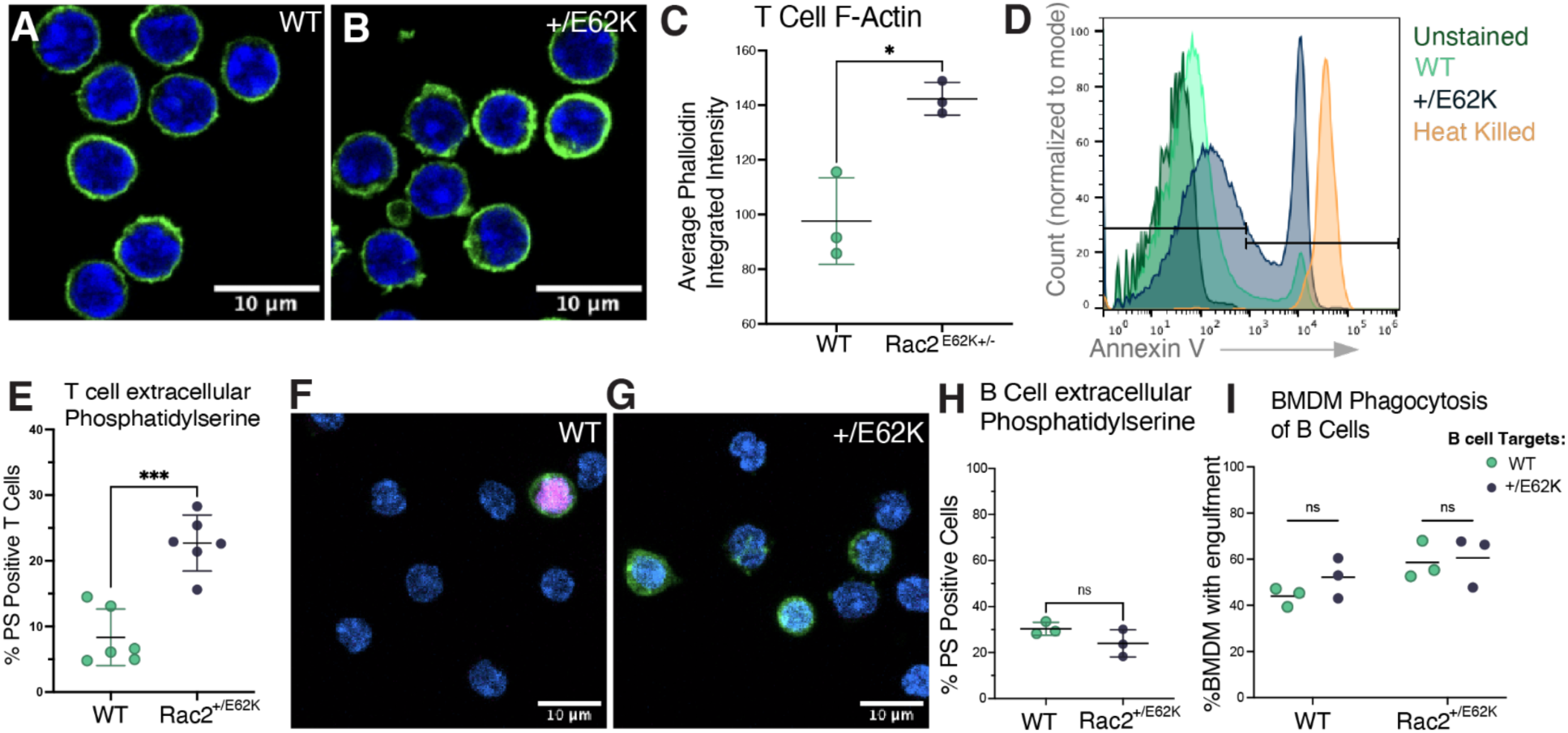
Rac2^+/E62K^ splenic T cells display elevated phosphatidylserine exposure consistent with enhanced phagocytic clearance. (A–B) Representative confocal images of WT (A) and Rac2^+/E62K^ (B) splenic T cells stained with phalloidin-488 (F-actin, green) and Hoechst (nuclei, blue). Scale bar = 10 µm. (C) Average F-actin phalloidin integrated density per mouse by Incucyte imaging. n=3 mice per genotype. Unpaired t-test. (D) Representative flow cytometry histogram of Annexin V binding on splenic T cells. Green = unstained; teal = WT; blue = Rac2^+/E62K^; orange = heat-killed positive control. The horizontal line indicates the PS-positive gate. (E) Percent phosphatidylserine-positive (Annexin V⁺) splenic T cells by flow cytometry. n=6 mice per genotype. Unpaired t-test. (F–G) Representative confocal images of WT (F) and Rac2^+/E62K^ (G) splenic T cells stained for extracellular phosphatidylserine (Annexin V, green), cell death (PI, magenta), and Hoechst (nuclei, blue). Scale bar = 10 µm. (H) Percent phosphatidylserine-positive (Annexin V⁺) splenic B cells by flow cytometry. n=3 mice per genotype. Unpaired t-test. (I) WT and Rac2^+/E62K^ BMDM phagocytosis of WT and Rac2^+/E62K^ splenic B cells measured by confocal imaging. n=3 mice per genotype. Two-way ANOVA with Tukey’s post-hoc test (factors: BMDM genotype, B cell genotype). Each dot represents one mouse. Data are mean ± SD; *p<0.05, **p<0.01, ***p<0.001, ****p<0.0001; ns = not significant.

Rac2^+/E62K^ bone marrow derived macrophages (BMDMs) exhibit enhanced phagocytosis of Rac2^+/E62K^ splenic T cells compared to wild type T cells (Mishra et al., 2023), suggesting phagocytic clearance as a possible mechanism for the profound T cell lymphopenia observed in human patients and in the mouse model. However, the ligand for macrophage recognition of T cells is unknown. Therefore, we measured phosphatidylserine (PS) exposure on T cells isolated from the spleen (representative histograms in Fig. 4D). Rac2^+/E62K^ T cells showed more than double the extracellular PS compared to wild type (Fig. 4E-G). Rac2^+/E62K^ T cells were also less viable over 20 hours of ex vivo culture (Fig. S4A). PS is a known “eat me” signal for phagocytes (Fadok et al., 1992). PS exposure therefore likely contributes to the enhanced phagocytosis of Rac2^+/E62K^ T cells by both wild type and Rac2^+/E62K^ macrophages.

In addition to the T cell lymphopenia, Rac2^+/E62K^ patients have a milder loss of circulating B cells (Hsu et al., 2019). We found similar levels of phosphatidylserine on the surfaces of splenic B cells in wild type and Rac2^+/E62K^ B cells. (Fig. 4H) and no significant difference in phagocytosis of either wild type or Rac2^+/E62K^ B cell genotype between wild type and Rac2^+/E62K^ BMDMs. This suggests that phagocytosis is not the driver for the B cell lymphopenia but is consistent with enhanced phagocytosis driving the T cell lymphopenia due to the combination of increased surface PS on Rac2^+/E62K^ T-cells and receptor-dependent hyperphagia of Rac2^+/E62K^ macrophages.

## Discussion

Mature Rac2^+/E62K^ neutrophils display hyperactivation markers including elevated F-actin, cell spreading, CD11b expression, and bioparticle internalization. The neutrophils also have reduced intracellular MPO and increased CD63 expression. The bone marrow development and architecture is normal, suggesting that reduced content reflects degranulation rather than a defect in granulopoiesis.

Rac2^+/E62K^ murine neutrophils have increased phagocytic uptake and intact phagosome acidification, but reduced MPO levels, which may be relevant to patient clinical phenotypes. RAC2^+/E62K^ patient neutrophils fail to prevent outgrowth of S. *aureus* (Smits et al., 2020). Efficient killing of S. *aureus* and *Aspergillus* depends on MPO-derived hypochlorous acid (HOCl) produced in the phagosome (Hampton et al., 1996). If Rac2^+/E62K^ neutrophil degranulation occurs prematurely, lower MPO availability could limit HOCl generation, severely limiting killing power. Although neutrophils have elevated upstream ROS production, this will not compensate for the MPO depletion (Odell and Segal, 1988). However, this theory needs to be tested by coculturing murine neutrophils with live S. *aureus* to understand whether the killing defect is recapitulated. Future experiments will investigate the disconnect between target internalization and killing.

Increased F-actin signal correlates with MPO reduction in Rac2^+/E62K^ neutrophils. Phalloidin intensity serves as a readout for Rac2 activity and provides a snapshot, but no information on the dynamic polymerization and turnover of actin filaments. Rac2 is a key selective regulator of primary granule exocytosis by neutrophils. Rac2 knockout has no effect on granule synthesis or density but abolishes primary granule exocytosis in a mouse model (Abdel-Latif et al., 2004). Stabilized cortical F-actin normally prevents premature granule fusion (Mitchell et al., 2008). Azurophilic degranulation requires RAC2-driven actin polymerization to move granules to the cortex, but coordinated local cortical actin depolymerization is necessary for granule fusion and exocytosis (Mitchell et al., 2008). RAC2 is best known for stimulating actin polymerization through Arp2/3, but in neutrophils specifically Rac2 signaling leads to the dephosphorylation of cofilin which activates it, stimulating F-actin severing (Sun et al., 2007).

This is in contrast to Rac activity in other cellular contexts where it activates LIM kinase to phosphorylate and inactivate cofilin. Thus Rac2 stimulates dynamic actin polymerization and turnover in neutrophils, which could be the mechanistic basis for the increased priming for degranulation.

Actin depolymerization is also required in NETosis (Thiam et al., 2020). RAC2^E62K^-expressing, dHL-60 neutrophil-like cells failed to undergo NETosis. MPO is required for PMA-induced NETosis in human (but not mouse) neutrophils (Metzler et al., 2011; Burn et al., 2025). MPO drives release of neutrophil elastase from primary granules which is required for chromatin decondensation. Although primary human neutrophils with partial MPO deficiency can still form NETs (Metzler et al., 2011), the combination of increased F-actin and decreased MPO may contribute to the NETosis defect in Rac2^+/E62K^ neutrophils because disassembly of the F-actin cytoskeleton is required during NETosis (Thiam et al., 2020).

The Rac2^+/E62K^ spleen is a myeloid-rich environment with T cells with elevated surface phosphatidylserine. This elevated PS is specific to Rac2^+/E62K^ T cells but not B cells, providing a plausible explanation for the disproportionately severe T cell lymphopenia in patients. We propose that this “eat me” signal could mediate the previously published BMDM hyper phagocytosis of T cells (Mishra et al., 2023). An interesting open question is the mechanism of PS exposure. PS exposure can arise from activation, exhaustion, or damage (Medina et al., 2026).

In this study, we show that Rac2^+/E62K^ neutrophils are hyperactivated and primed for azurophilic granule degranulation. MPO depletion provides a candidate mechanism for the inability of Rac2^+/E62K^ patient neutrophils to control S. *aureus* colonization despite increased phagocytic capacity, in addition to the previously described fMLF chemotaxis defect in patient blood neutrophils (Hsu et al., 2019; Smits et al., 2020). Splenic extramedullary granulopoiesis creates a myeloid-rich environment in which neutrophils with elevated oxidative burst and degranulation as well as hyperphagocytic monocytes accumulate. T cells exhibit elevated phosphatidylserine exposure, an eat-me signal that sensitizes them to phagocyte-mediated clearance. Together these findings suggest that the Rac2^+/E62K^ mutation may compromise antimicrobial function in neutrophils and drive selective T cell clearance in the spleen, both of which likely contribute to the recurrent infections experienced by human patients.

## Methods

### Mouse Husbandry

Mice were maintained in sterile living conditions at the Animal Resource Center, an AAALAC accredited animal facility, at The University of California, Santa Barbara (UCSB). Rac2+/E62K mice were kindly provided by Amy Hsu (National Institute of Allergy and Infectious Diseases (NIAID), National Institutes of Health) and backcrossed to C57BL/6 on site. Procedures were performed in accordance with Institutional Animal Care and Use Committee (IACUC) guidelines and approved protocols. Ear punch biopsies were used for genotyping by Transnetyx, an automated genotyping service. 8-12-week-old female and male mice were used for all experiments.

### Histology

Femurs and spleens were harvested from 8-12-week-old mice and fixed in 4% PFA over night. Fixed tissues were submitted to IDEXX BioAnalytics for paraffin embedding, sectioning, hematoxylin and eosin (H&E) staining and mounting. Femurs were imaged by IDEXX at 1X and 40X magnification. Representative H&E spleen images were taken at 5X and 10X magnification using an Axiovert 40 CXL. Tissues were assessed by a board-certified veterinary pathologist (IDEXX BioAnalytics).

### Spleen Weights

Spleens were isolated from 8-12 week old mice, trimmed of adherent fat and connective tissue, and blotted dry on absorbent paper prior to weighing. Organ mass was recorded to the nearest 0.1 mg using a calibrated analytical balance (Mettler Toledo XPR205, Columbus, OH).

### Cell Isolation

#### Bone Marrow Neutrophil Isolation

Femurs and tibiae were harvested from 8-12-week-old mice. Bone marrow was flushed with FACS buffer (PBS supplemented with 2% FBS and 1mM EDTA) or mechanically dissociated and filtered through a 40um cell strainer (Corning Falcon #08-771-1). Neutrophils were isolated using the EasySep™ Mouse Neutrophil Enrichment Kit (STEMCELL Technologies #19762) per the manufacturer protocol. Enriched neutrophils were resuspended in lymphocyte medium (RPMI 1640 supplemented with 10% heat-inactivated FBS and 1% penicillin/streptomycin), kept on ice, and used within 1 hour. Purity assessed by Ly6G and CD11b staining and flow cytometry.

#### Splenic cell Isolation (Neutrophils, T cells, B cells)

Spleens were harvested from 8-12-week-old mice dissociated through a 70µm cell strainer (Corning Falcon #352350) and neutrophils isolated with EasySep™ Mouse Neutrophil Enrichment Kit (STEMCELL Technologies #19762) per the manufacturer protocol. T cells were isolated with EasySep™ Mouse T Cell Isolation Kit (STEMCELL Technologies #19851) per manufacturer protocol. For B cell assays, the EasySep Mouse B Cell Isolation Kit (STEMCELL Technologies #19854) was used per the manufacturer protocol. All splenic cells were isolated using EasyEights™ EasySep™ Magnet (Stemcell Technologies, #18103).

#### Single-Cell Suspension (Immunophenotyping)

Cells were centrifuged at 400xg for 4 minutes at 4 degrees C. Red blood cells were lysed in 1 mL 1X RBC Lysis Buffer (Invitrogen™ 50-112-9751) for 3 minutes (bone marrow) or 4 minutes (spleen) at room temperature, quenched with 10 mL ice cold FACS buffer, and centrifuged at 400xg for 4 minutes at 4 degrees C. Cells were resuspended in 5mL of FACS buffer, kept on ice, and counted.

### Flow Cytometry Immunophenotyping

For immunophenotyping, whole bone marrow or whole spleen single-cell suspensions were prepared as described above following RBC lysis. 1 × 10^6 cells per well were plated in a 96 Well Conical (V) Bottom Plate (#277143) in 150 µL PBS. Live/Dead discrimination used the LIVE/DEAD Fixable Violet Dead Cell Stain Kit (Thermo Fisher #L34963; 1:100 in protein-free PBS, 20 minutes at 4 degrees C in the dark), followed by 2X washing with 200 µL PBS. Cells were resuspended in 150uL of FACS buffer, and Fc receptors were blocked with 25 µL Purified anti-mouse CD16/32 antibody (BioLegend #101301; 1:100 final in FACS buffer) for 10 minutes at 4 degrees C in the dark without washing. 25uL of antibody mastermix was then added directly to wells and incubated for 20-30 minutes at 4 degrees C in the dark. Cells were washed twice with 200 µL FACS buffer. Cells were further diluted with 1mL of FACS buffer and run on an Attune NxT flow cytometer. Data were analyzed using FlowJo (version 10).

Ultra comp beads were used for compensation. Fluorescence minus one controls were used to set gates. We followed the gating strategy outlined in Evrard et al. (2018). The complete antibody panel is shown below.

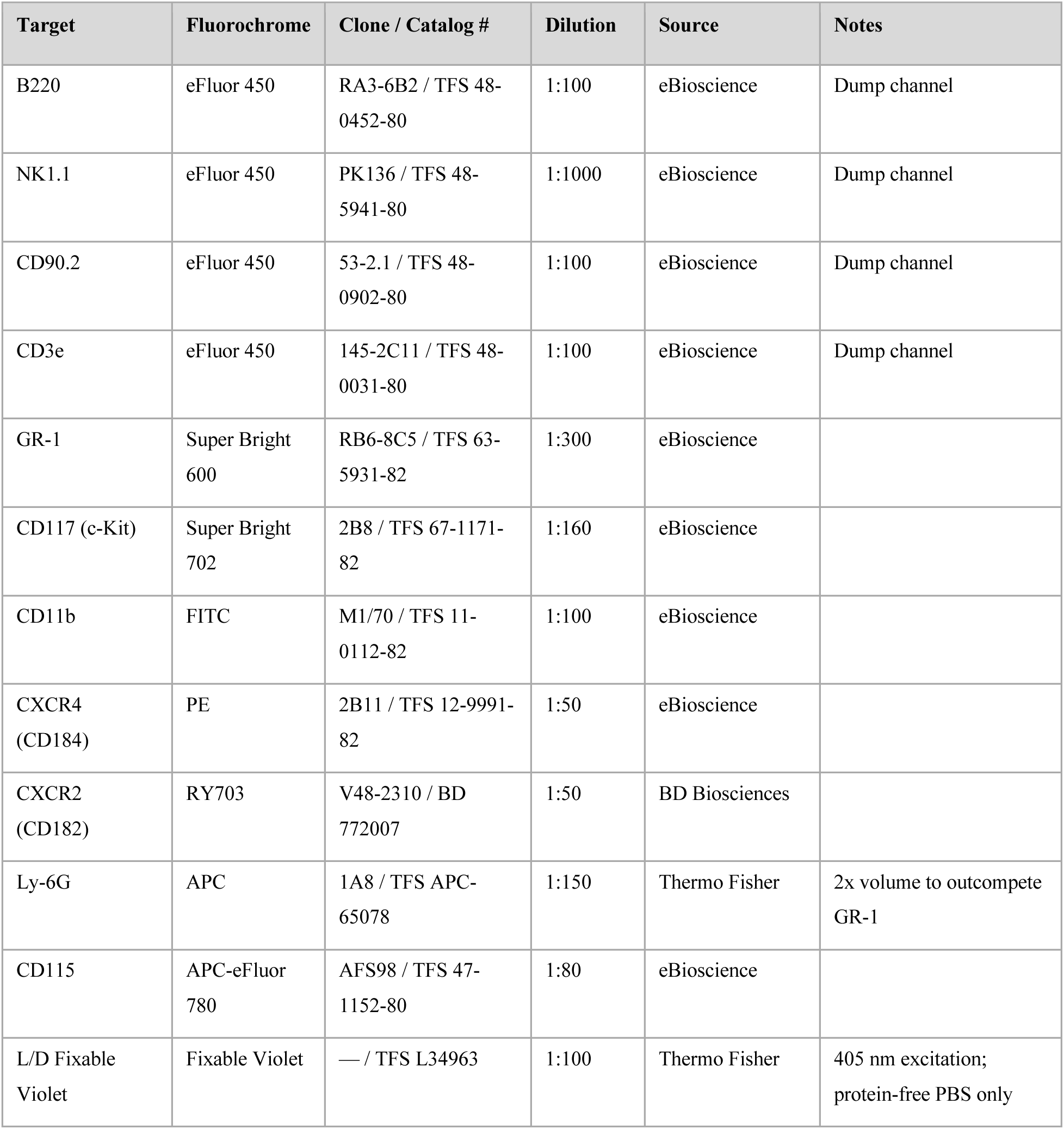

| Target | Fluorochrome | Clone / Catalog # | Dilution | Source | Notes |
| --- | --- | --- | --- | --- | --- |
| B220 | eFluor 450 | RA3-6B2 / TFS 48-0452-80 | 1:100 | eBioscience | Dump channel |
| NK1.1 | eFluor 450 | PK136 / TFS 48-5941-80 | 1:1000 | eBioscience | Dump channel |
| CD90.2 | eFluor 450 | 53-2.1 / TFS 48-0902-80 | 1:100 | eBioscience | Dump channel |
| CD3e | eFluor 450 | 145-2C11 / TFS 48-0031-80 | 1:100 | eBioscience | Dump channel |
| GR-1 | Super Bright 600 | RB6-8C5 / TFS 63-5931-82 | 1:300 | eBioscience |  |
| CD117 (c-Kit) | Super Bright 702 | 2B8 / TFS 67-1171-82 | 1:160 | eBioscience |  |
| CD11b | FITC | M1/70 / TFS 11-0112-82 | 1:100 | eBioscience |  |
| CXCR4 (CD184) | PE | 2B11 / TFS 12-9991-82 | 1:50 | eBioscience |  |
| CXCR2 (CD182) | RY703 | V48-2310 / BD 772007 | 1:50 | BD Biosciences |  |
| Ly-6G | APC | 1A8 / TFS APC-65078 | 1:150 | Thermo Fisher | 2x volume to outcompete GR-1 |
| CD115 | APC-eFluor 780 | AFS98 / TFS 47-1152-80 | 1:80 | eBioscience |  |
| L/D Fixable Violet | Fixable Violet | — / TFS L34963 | 1:100 | Thermo Fisher | 405 nm excitation; protein-free PBS only |

### Intracellular Myeloperoxidase Staining (Primary Neutrophils)

1E6 EasySep-enriched bone marrow neutrophils were collected, centrifuged at 500xg for 5 minutes at room temperature, and resuspended in 400 µL 1X IC Fixation Buffer (eBioscience #00-8222-49) for 20 minutes at 4 degrees C. Cells were centrifuged at 600xg for 5 minutes at 4 degrees C and resuspended in 500uL 1X Perm Buffer (10x Perm concentrate diluted 1:10 in ultrapure H_2_O (eBioscience #00-8333). Cells were permeabilized for 20 minutes at 4C then centrifuged at 600xg for 5 minutes at 4C. Cells were washed with FACs buffer. Cells were then incubated with anti-MPO polyclonal primary antibody (Thermo Fisher #PA5-16672; 1:100) in BlockAid Blocking Solution (Invitrogen #B10710) supplemented with mouse buffer and EDTA (100 µL per well) overnight at 4 degrees C. After washing in 1X Perm buffer, goat anti-rabbit Alexa Fluor 647 secondary antibody (Invitrogen #A21245, highly cross-adsorbed H+L; 1:100), (Hoechst 1:500), and (Phalloidin 488 1:500) was applied for 30 minutes at room temperature in the dark. Cells were washed 2X in FACS buffer and finally resuspend in PBS and plated on a Poly-L-Lysine (Catalog#?) coated 96well glass bottom plate for imaging (Catalog#?). Images were acquired on a Zeiss LSM980 confocal microscope at 63x oil immersion objective using laser lines 405 nm (Hoechst), 488 nm (Alexa Fluor 488 Phalloidin), and 647 nm (Alexa Fluor 647).

### Quantitative Image Analysis For MPO levels

#### Cellpose segmentation

Cell segmentation was performed using CP-SAM (Cellpose Segment Anything Model) in Cellpose (version 4.0.6). CP-SAM was run with a diameter parameter of 150 pixels.

Segmentation masks were created from a single z slice. They were manually inspected and corrected when necessary to ensure accurate cell boundaries.

#### Fiji Analysis

Image analysis was performed using a custom batch-processing macro in Fiji (Image J version 2.16.0/1.54p). To perform cell-by-cell quantification, segmentation masks generated in Cellpose were automatically matched to corresponding fluorescence images. All analysis was run on the same image slice the masks were created from. The macro converted each labeled cell from the segmentation mask into regions of interest (ROIs) that were applied to all three fluorescent channels. Mean fluorescence intensity and integrated density measurements were collected for MPO, F-actin, and nuclear signals in each cell. Cell area measurements were also collected. Any cells touching image borders were automatically excluded from analysis. No background subtraction was performed. For batch normalization, each individual cell’s MPO IntDen was divided by the mean WT IntDen from the same imaging session, yielding a per-cell normalized value (MPO_norm) with the WT mean set to 1.0 per experiment. For the F-actin quartile analysis, normalized F-actin IntDen values were pooled across all sessions and divided into four equal quartiles (Low, Med-Low, Med-High, High). Each cell was assigned to a quartile bin based on its individual normalized F-actin value. Per-mouse mean MPO_norm was then calculated within each bin. The statistical tests are run on per-mouse means.

### T cell F-Actin measurements

EasySep-enriched splenic T cells were fixed in 400 µL IC Fixation Buffer for 20 minutes at 4 degrees C. After washing with 3 mL FACS buffer and centrifugation at 600xg for 5 minutes at 4 degrees C, cells were permeabilized in 650 µL 1x Permeabilization Buffer. 100 µL was transferred per well, centrifuged, and cells stained with Alexa Fluor 488 Phalloidin (Invitrogen #A12379; 1:500; 2 µL stock per 1000 µL FACS buffer, 100 µL per well) for 20 minutes at 4 degrees C in the dark. After two PBS washes, cells were imaged on the Incucyte Zoom (Sartorius) at 20x magnification using non-adherent cell analysis.

### Cell Spreading

Glass-bottom 96-well plates were coated with fibronectin (Sigma #F1141, 1:100 in PBS, 60 µL per well, 1 hour at room temperature). 50,000 EasySep-enriched bone marrow neutrophils per well were plated in 100 µL complete RPMI with or without 5 ng/mL GM-CSF (PeproTech #315-03) and incubated at 37 degrees C, 5% CO_2_. Cells were imaged at 2 hours on the Incucyte Zoom (Sartorius) at 20x magnification. Spreading area per cell was quantified using Incucyte adherent Cell-By-Cell analysis software. Statistical comparison by two-way ANOVA with Tukey post-hoc test (factors: genotype, GM-CSF stimulation).

### NETosis Assay

HL-60 cells used in this study were obtained from Orion Weiner’s lab at UCSF. Cells were maintained at 37 °C with 5% CO_2_ in RPMI 1640 medium (Gibco, # 22400089) with 10% heat inactivated fetal bovine serum (Sigma-Aldrich #F4135). dHL-60 neutrophil-like cells were differentiated from HL-60 in 1.25% DMSO and 10% FBS for 5 days. On day 5, GFP or RAC2E62K expression was induced with doxycycline hydrochloride (final: 1ug/mL ; #J67043-ADJ67043), in 1.25% DMSO in complete RPMI . On day 6, Poly-L-Lysine-coated 96-well plates (coated overnight, washed with H_2_O and dried) were seeded with 20,000 dHL-60 cells per well in 100 µL RPMI supplemented with no FBS. PMA (#tlrl-pma; Fisher Scientific) was added at final concentrations of 0, 25, 50, or 100 nM to a final volume of 200uL. Cells were incubated at 37 degrees C, 5% CO_2_ for five hours. SYTOX Orange Nucleic Acid Stain (5 mM in DMSO; Thermo Fisher #S11368; 1:1000 final dilution, 50 µL per well) was added. Plates were centrifuged at 400xg for 5 minutes at 4 degrees C and 150 µL supernatant carefully removed. Fluorescence was measured on a Molecular Devices SpectraMax iD3 plate reader (excitation 546 nm, emission 570 nm).

### dHL-60 Intracellular MPO Staining

250,000 cells per well were collected, centrifuged at 500xg for 5 minutes at room temperature, and resuspended in 400 µL IC Fixation Buffer (1x) for 20 minutes at 4 degrees C. Cells were centrifuged at 600xg for 5 minutes at 4 degrees C and washed twice with 1x Permeabilization Buffer (10x PERM concentrate diluted 1:10 in ultrapure H_2_O, 200 µL per wash). Cells were resuspended in anti-MPO-PE antibody (clone MPO455-8E6; Thermo Fisher #12-1299-42) at 1:50 in 100 µL FACS buffer and incubated for 40 minutes at 4 degrees C. After two washes, cells were resuspended in 200 µL FACS buffer and acquired on the Attune NxT.

### Transwell Migration Assay

Bone marrow neutrophil chemotaxis was assessed using Corning Transwell polycarbonate membrane inserts (6.5 mm diameter, 5.0 µm pore; Sigma-Aldrich #CLS3421-48EA). 200,000 EasySep-enriched neutrophils in 200 µL lymphocyte medium were placed in the upper chamber. The lower chamber contained 600 µL lymphocyte medium with 0 or 20 ng/mL CXCL2 (PeproTech #250-15). Plates were incubated at 37 degrees C, 5% CO_2_ for 1 hour. Migrated cells were quantified using the Incucyte Zoom (Sartorius) non-adherent cell analysis module.

### Phosphatidylserine Exposure Measurements

EasySep-enriched splenic T and B cells were isolated as described above. Staining used the BioLegend FITC Annexin V Apoptosis Detection Kit with PI (#640914) per the manufacturer protocol. 5 × 10^6 cells per sample were pelleted, resuspended in Annexin V binding buffer, incubated with FITC-Annexin V and propidium iodide for 15 minutes at room temperature in the dark, diluted with binding buffer, and acquired immediately on the Attune NxT. Heat-killed cells served as the positive control for gate setting. For confocal imaging of extracellular PS, cells were stained with fluorescently conjugated Annexin V (green), PI (cell death, magenta), and Hoechst 33342 (Invitrogen #H1399; nuclei, blue) and imaged on the Zeiss LSM800 at 40x.

### pHrodo Bioparticle Phagocytosis Assay

Glass-bottom 96-well plates (Cellvis #P96-1.5H-N) were fibronectin-coated (1:100) (Sigma #F1141) and incubated at 37C for 3 hours before removing. 50,000 EasySep-enriched bone marrow neutrophils per well were plated in 100 µL complete RPMI and allowed to adhere for approximately 40 minutes in the Incucyte Zoom at 37 degrees C, 5% CO_2_. pHrodo Red bioparticles: E. coli (Thermo Fisher #P35361), Zymosan A (Thermo Fisher #P35364), and S. aureus (Thermo Fisher #A10010), were resuspended in 2mL of PBS and sonicated for 10 minutes. Bioparticles were added at 10 µg/well for E. *coli* and S *aureus*, and 5ug/well for Zymosan in 50 µL per well. Live-cell imaging proceeded at 20-minute intervals for 5 hours at 37 degrees C, 5% CO_2_ on the Incucyte Zoom at 20x magnification. Total integrated pHrodo intensity per cell and percentage of pHrodo-positive cells were quantified using Incucyte Cell-By-Cell analysis software.

### Preparation and culture of bone marrow-derived macrophages

Femurs and tibias were harvested from 8-12-week-old mice and crushed to isolate whole bone marrow (WBM) cells. WBM cells were filtered through 40µm nylon cell strainers and incubated with 1X RBC Lysis solution (Thermo, #50-112-9751) to eliminate red blood cells. Cells were plated with L-929 conditioned media and differentiated for seven days to produce bone marrow-derived macrophages (BMDMs).

### L929-conditioned media (BMDM media)

To generate L929-conditioned media, L929 cells were cultured until confluent then passaged and grown for 10 days. On day 10, the media containing M-CSF was collected and filtered through a 40µm nylon cell strainer and aliquots were stored in -20oC. BMDMs were subsequently grown in BMDM media: RPMI Glutamax (Gibco, #72400047), 20% MCSF L929-conditioned media, 10% heat-inactivated FBS and 1% Pen-strep.

### Bone Marrow Derived Macrophage: B cell Phagocytosis assay

75,000 BMDMs were stimulated with IFN-γ (Biolegend, #50-170-393) for 4hrs on day seven and plated in BMDM media in a 96-well glass bottom plate (Cellvis, #P96-1.5H-N) overnight. On day eight, BMDM media was changed to mcsf-free media prior to the experiment. After isolation, B cells were stained with CellTrace Far Red (Thermo, #C34564) and pHrodo Red SE (Invitrogen #P36600). 375,000 B cells were added to the wells and co-cultured with the BMDMs overnight. Next day, samples were fixed with 4% PFA and permeabilized with PBS-T (0.1% Triton-X). BMDMs were stained with anti-F4/80 (Abcam, #ab6640) and IgG (H+L) Cross-Adsorbed Goat anti-Rat, Alexa Fluor® 488 (Thermo, #A-11008). DAPI was used to visualize DNA. Engulfment was quantified as a CellTrace Far Red or pHrodo signal within a (green) macrophage. Data was acquired on the Leica DMi8 Epifluorescence scope and processed on ImageJ/Fiji for quantification.

### Statistical Analysis

Statistical analyses were performed in GraphPad Prism or Python. Statistical significance is indicated as ∗p<0.05, ∗∗p<0.01, ∗∗∗p<0.001, ∗∗∗∗p<0.0001. Error bars represent mean ± SD unless otherwise noted. ns = not significant.

### Use of AI

During the preparation of this manuscript, the authors used Claude (Anthropic) to assist with editing and organization of writing and to assist with exploratory data analysis approaches, including the selection of statistical tests and binning strategies for the F-actin/MPO quartile analysis (Fig. 1F). All AI-generated content was reviewed, edited, and verified by the authors. AI was not used to draft scientific conclusions.

