## Supplemental Files for "Rac2 Hyperactivity Drives Neutrophil Degranulation, Myeloperoxidase Deficiency, and Lymphopenia"

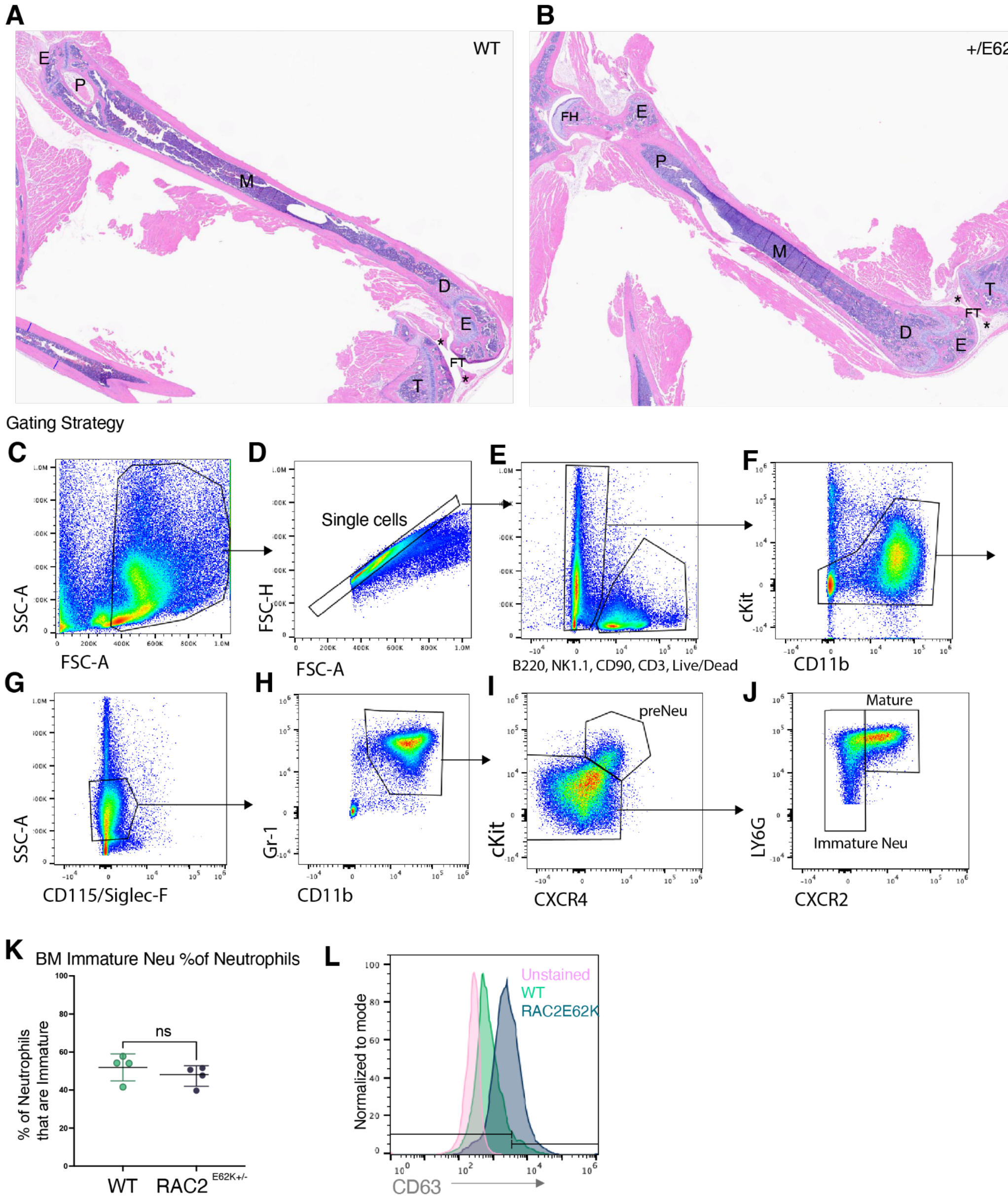


**Supplemental Figure 1.** (A–B) Hematoxylin and eosin stained femur sections from WT (A) and RAC2$^{+/E62K}$ (B) mice at 1× magnification. Orientation differs slightly between samples, but proximal (P), distal (D), and mid (M) diaphysis and epiphyses (E) are visible in both. The proximal tibia (T), portions of menisci (*), femorotibial joint (FT), and femoral head (FH) sitting in the acetabulum of the pelvis are also seen. No differences in bone or bone marrow morphology were identified by board-certified veterinary pathologist review (IDEXX BioAnalytics). (C–J) Representative flow cytometry gating strategy used to identify neutrophil developmental populations in bone marrow, adapted from Evrard et al. (2018). (C) Cells gated by FSC-A vs SSC-A. (D) Singlets gated on FSC-A vs FSC-H. (E) Lineage-negative live cells identified by exclusion of B220, NK1.1, CD90, CD3, and a viability dye. (F) cKit vs CD11b gating to separate cKit$^{+}$CD11b$^{-}$ progenitors from cKit$^{-}$CD11b$^{+}$ committed cells. (G) Exclusion of CD115$^{+}$/Siglec-F$^{+}$ monocytes and eosinophils. (H) Gr-1 vs CD11b gating to identify the Gr-1$^{+}$CD11b$^{+}$ neutrophil lineage population. (I) Pre-neutrophils (preNeu) identified as cKit$^{+}$CXCR4$^{+}$. (J) Mature (Ly6G$^{+}$CXCR2$^{+}$) and immature (Ly6G$^{+}$CXCR2$^{-}$) neutrophils gated from the cKit$^{-}$ population. (K) Immature neutrophils as a percentage of total bone marrow neutrophils by flow cytometry. n=4 mice per genotype. Unpaired t-test. Each dot represents one mouse. (L) Representative flow cytometry histogram of CD63 surface expression on mature bone marrow neutrophils. Pink = unstained; teal = WT; blue = Rac2$^{+}$^/E62K. The horizontal line indicates the CD63-positive gate. Data are mean ± SD; *$p<0.05$, **$p<0.01$, ***$p<0.001$, ****$p<0.0001$; ns = not significant.

### Supplemental 2

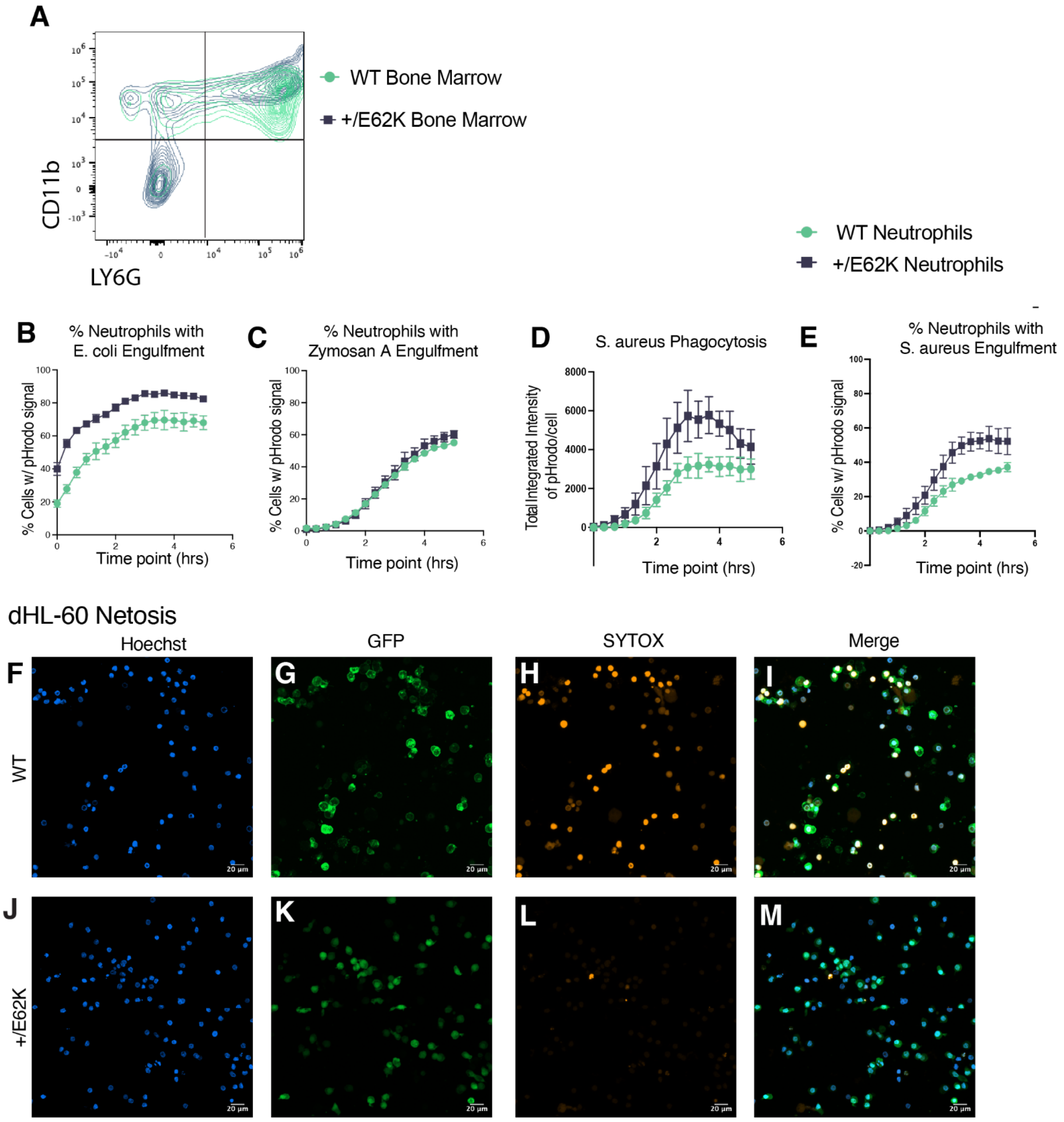

### Supplemental 3

**A** Mouse Spleens

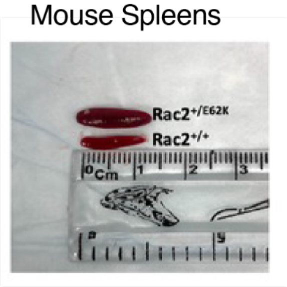


### Supplemental 4

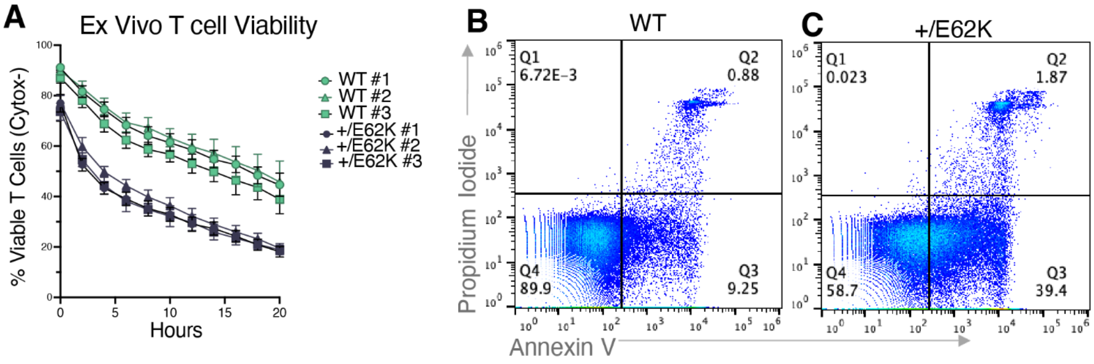
